# Endosperm development is an autonomously programmed process independent of embryogenesis

**DOI:** 10.1101/2020.08.31.275354

**Authors:** Hanxian Xiong, Wei Wang, Meng-Xiang Sun

## Abstract

The seeds of land plants contain three genetically distinct structures: the embryo, endosperm, and seed coat. The embryo and endosperm need to interact and exchange signals to ensure coordinated growth. Accumulating evidence has confirmed that embryo growth is supported by the nourishing endosperm and regulated by signals originating from the endosperm. Available data also support that endosperm development requires communication with the embryo. Here, using single-fertilization mutants, *Arabidopsis dmp8/9* and *gex2*, we demonstrate that in the absence of a zygote and embryo, endosperm initiation, syncytium formation, free nuclear cellularization, and endosperm degeneration are as normal as in the wild type in terms of the cytological process and time course. Although rapid embryo expansion accelerates endosperm breakdown, our findings strongly suggest that endosperm development is an autonomously organized process, independent of egg cell fertilization and embryo–endosperm communication. This work confirms both the altruistic and self-directed nature of the endosperm during coordinated embryo-endosperm development. The findings provide novel insights into the intricate interaction between the two fertilization products and will help to distinguish the real roles of the signaling between endosperm and embryo. These finding also shed new light on agro-biotechnology for crop improvement.

## INTRODUCTION

Seed development in flowering plants is initiated by double fertilization, which leads to the formation of a diploid zygotic embryo and triploid endosperm. These two genetically distinct “siblings” then develop concomitantly within the surrounding maternal tissues, the seed coat, to form a seed (Lafon-Placette and Kohler, 2014). The endosperm plays an important role in supporting embryo growth by supplying nutrients and other factors during seed development and germination (Ingram, 2020; Li and Berger, 2012). Several endosperm-expressed genes, such as *EMBRYO SURROUNDING FACTOR 1* (*ESF1*), *ABNORMAL LEAF SHAPE1* (*ALE1*), and *ZHOUPI* (*ZOU*) (Costa et al., 2014; Tanaka et al., 2001; Yang et al., 2008) have been reported to regulate embryo development. Endosperm cellularization also defines an important developmental transition for embryo development (Hehenberger et al., 2012). In mutants of *fertilization independent seed 2* (*fis2*) and *endosperm defective 1* (*ede1*) that fail to undergo endosperm cellularization, embryo development is arrested. A recent work first clearly describes a pathway for the communication between the endosperm and embryo, in which *TWISTED SEED1* (*TWS1*) acts as a ligand of the receptor-like kinases GSO1 and GSO2 in the embryo and this sulfated peptide needs to be cleaved by ALE1 in the neighboring endosperm to release the active peptide, which then triggers GSO1/2-dependent cuticle reinforcement in the embryo (Doll et al., 2020). This strongly suggests that normal endosperm is essential for embryo development. Conversely, some embryo-derived factors have also been reported to regulate endosperm development, reflecting the impact of embryo development on endosperm (Aw et al., 2010; Nowack et al., 2006; Xu et al., 2015). Mutant analysis using defective kernel mutations in maize also provided examples showing that the normal embryo could enhance the mutant endosperm development (Neufferand Sheridan, 1980). It sees gradually accepted that embryo and endosperm development depend on each other. Since embryogenesis and endosperm development are also involved in the seed coat development, it is critical to further clarify the relationship among them and especially between the two fertilization products to recognize the roles of cell-cell communications in their coordinated development.

Two membrane proteins, GAMETE-EXPRESSED 2 (GEX2) (Mori et al., 2014) and DOMAIN OF UNKNOWN FUNCTION 679 membrane protein 8/9 (DMP8/9) (Cyprys et al., 2019; Takahashi et al., 2018), have been reported to be involved in double fertilization and loss of their functions caused single central cell fertilization, offering a unique opportunity to exclude an embryonic effect on endosperm development and to investigate the need of an embryo for endosperm development at every critical stage, as well as the seed coat development in an embryo-free seed.

## RESULTS AND DISCUSSION

To address this, we first created *AtDMP8*/*9* CRISPR/Cas9 gene-edited double mutants, and found that these mutants showed serious defects in the seed set, similar to those previously reported (Figures 1A to 1F) (Cyprys et al., 2019). Obviously, knockout of *DMP8*/*9* could produce embryo-free seeds by single fertilization (Figures 1G to 1J). Thus, based on the mutants, we then characterized the main features of early endosperm development by clearing seeds at successive development stages when the embryo was not present (Figure 2). We found that when only the central cell was fertilized, the primary endosperm cell divided normally, indicating that the initiation of endosperm development does not require a message from the zygote and an unfertilized egg cell does not negatively affect endosperm development. After the first divisions of the primary endosperm nucleus, one nucleus migrated along the micropylar-chalazal axis to near the unfertilized egg cell, just like its counterpart in Col-0 ovules moves toward the zygote. After the third nuclear division, one or two nuclei were located at the chalazal pole of the embryo sac, leading to eight endosperm nuclei evenly distributed along a curved tube-like embryo sac. During the following cycles of syncytial division, the larger nuclei were observed in the posterior endosperm pole, which is an early marker of the chalazal endosperm. Then, the syncytial endosperm continued to divide until the endosperm nuclei fully distributed in the periphery rejoin of the embryo sac. These results confirm that the embryo-free seeds undergo the same endosperm development pattern and follow the same time course as the wild type (WT) in the syncytial phase, in terms of endosperm initiation, nuclear migration, and free nucleus distribution (Figure 2; Supplemental Figure 1A). In addition, the Green fluorescent protein (GFP) reporters of four endosperm marker genes (Li et al., 2010; Portereiko et al., 2006; Steffen et al., 2007) were expressed in embryo-free seeds (Figures 3A to 3D) and the micropylar endosperm, which occupies a domain called the embryo-surrounding region (ESR), was also marked by the basic helix loop helix factor-*ZHOUPI* (*ZOU*) (Yang et al., 2008), and its expression pattern and levels in embryo-free seeds remained normal, as in the WT (Figures 3E and 3F). These observations suggest that the formation or presence of an embryo is not required for early syncytial endosperm development.

**Figure 1.**
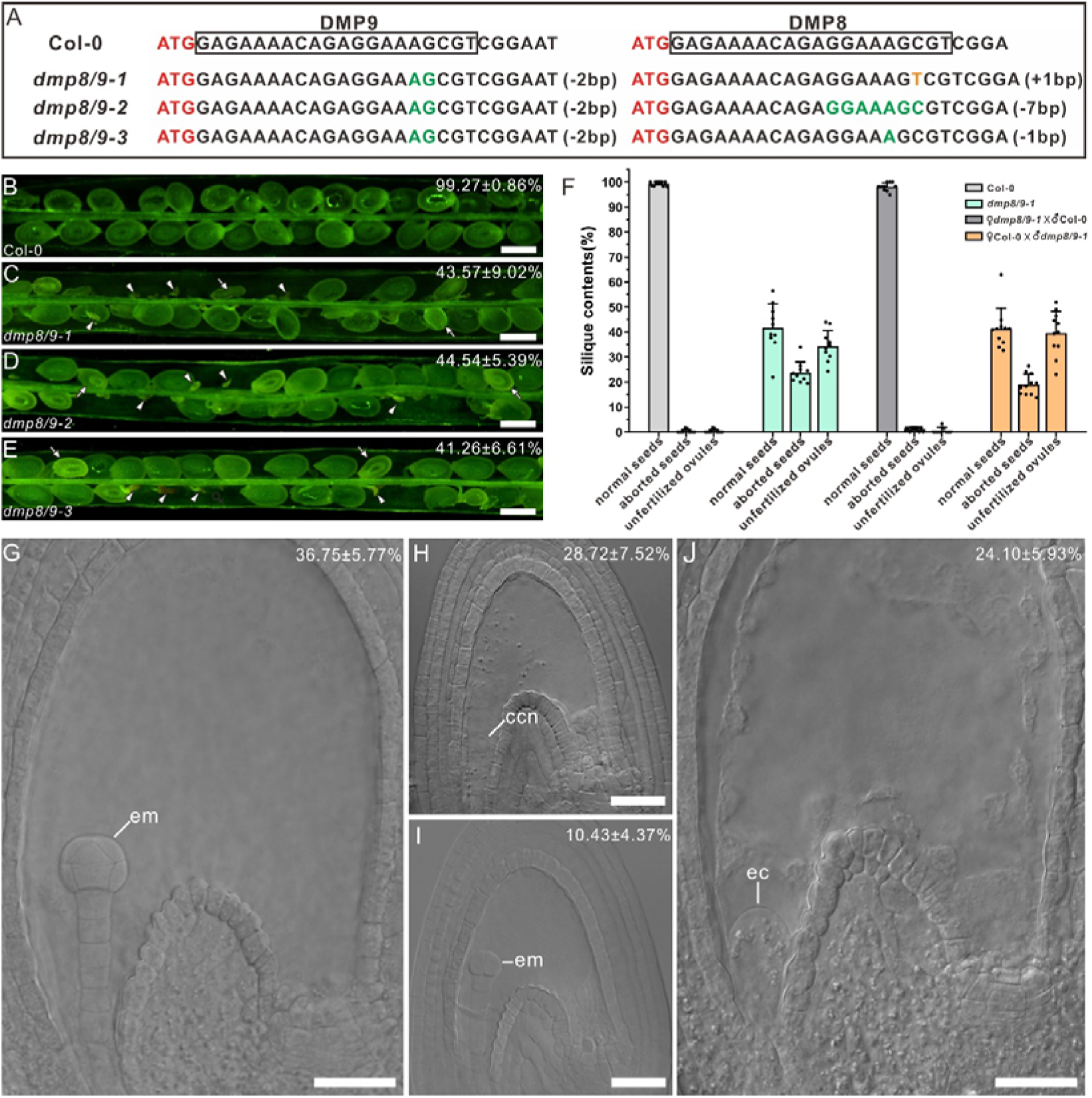
Knock-out of *DMP8* and *DMP9* causes single-fertilization. **(A)** Targeted mutagenesis in *Arabidopsis* using the CRISPR/Cas9 system by one target site. The black boxes showed the target site and disrupted *DMP8* and *DMP9* coding sequences just after ATG in three different homozygous mutants. **(B-E)** Unfertilized ovules (arrowheads) and aborted seeds (arrows) were frequently observed in three *dmp8/9* double mutants generated by CRISPR/Cas9. The seed set rates (means ± SD) were shown at the top right. n=568, 723, 539 and 467 seeds, respectively. Scale bar represents 0.5mm. **(F)** Statistics of various types of seed abortion in different crossing groups for Col-0 and *dmp8/9-1*. For each crossing group, from left to right, n=631, 551, 610 and 627 seeds, respectively. Data are the means ± SD. **(G-J)** DMP8/9 are required for fertilization. The phenotype of non-fertilization **(H)** or single-fertilization **(I** and **J)** were observed in *dmp8/9-1* seeds (n=391). The data (means ± SD) were shown at the top right. Scale bars=20µm. Abbreviations: em, embryo; ec, the egg cell; ccn, the central cell nuclei.

**Figure 2.**
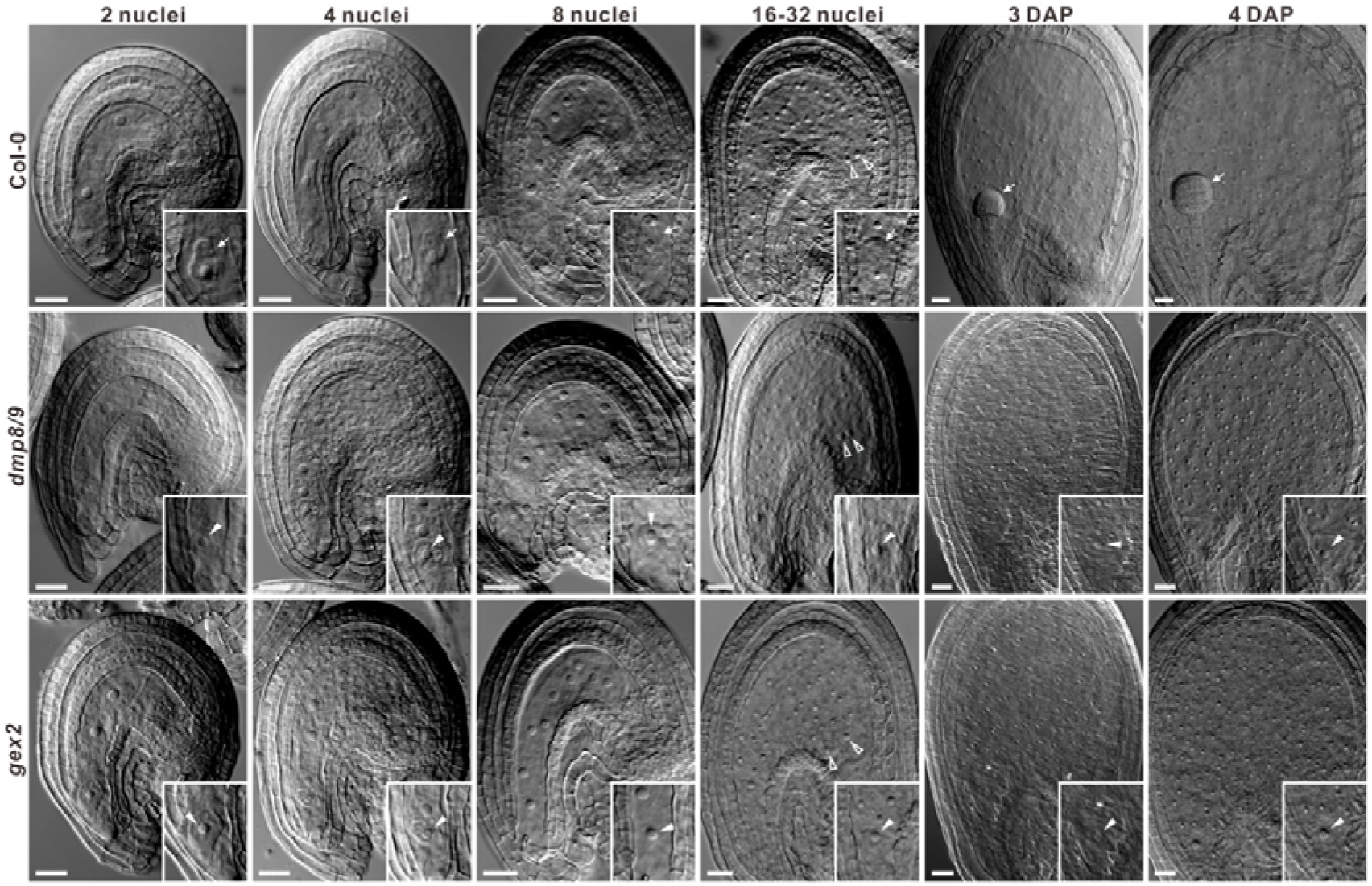
Early endosperm development does not depend on the presence of an embryo. Cleared seeds before 4 DAP. The ovules were pollinated with Col-0, *dmp8/9*, or *gex2* pollen. Note that when only the central cell was fertilized, endosperm development initiates and proceeds normally, just as in Col-0 seeds. The insets in the lower right indicate egg cell nuclei (arrows), zygotes or embryos (arrowheads). Hollow arrowheads indicate larger nuclei in the chalazal endosperm. Scale bars, 20 µm.

**Figure 3.**
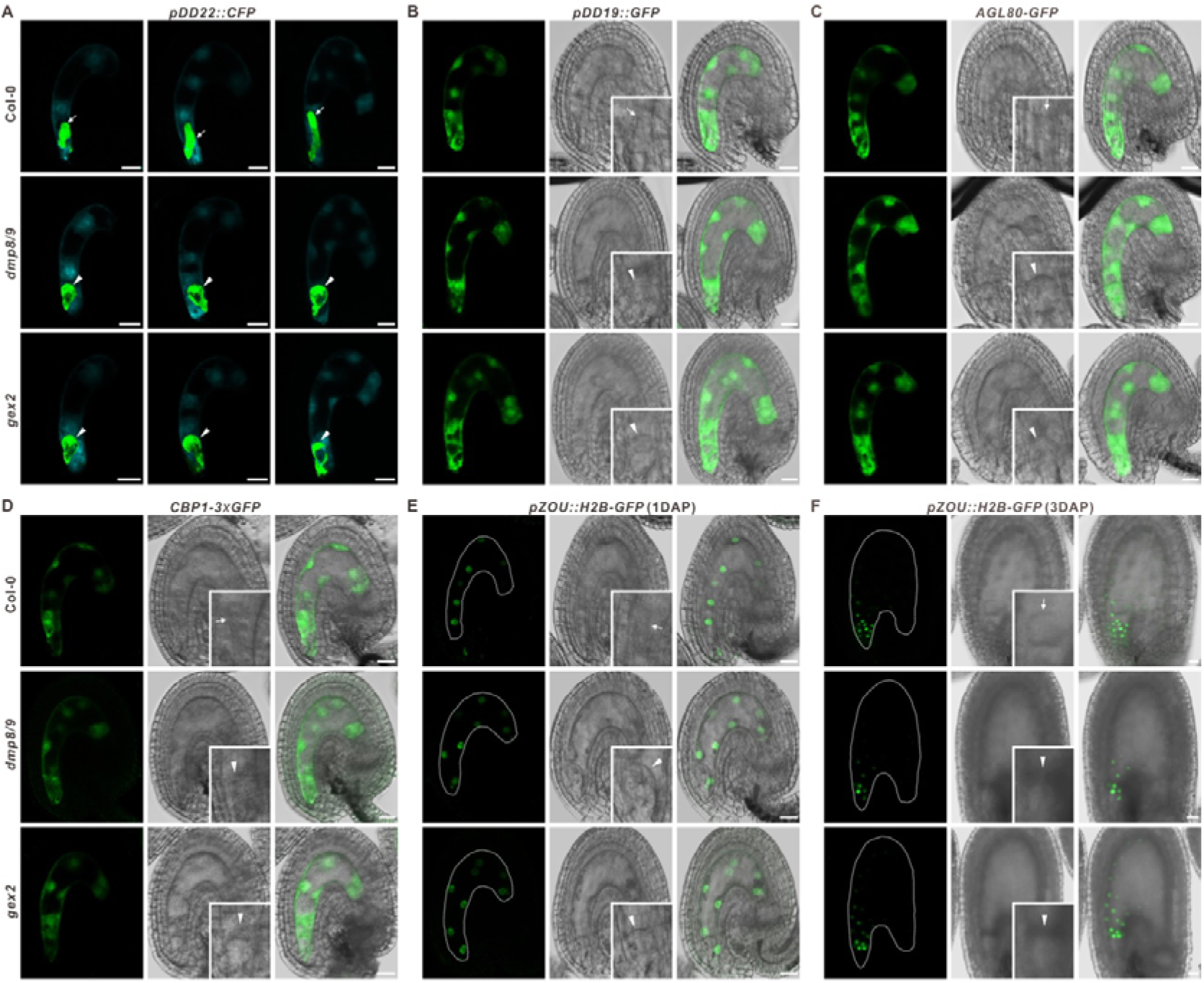
Embryo-free seeds express different endosperm-cell fate markers. **(A–F)** Confocal laser scanning microscopy (CLSM) images of seeds expressing different endosperm reporters: *pDD22::CFP* in **(A)**, *pDD19::GFP* in **(B)**, *pAGL80::AGL80-GFP* in **(C)**, *pCBP1::CBP1-3xGFP* in **(D)**, and *pZOU::H2B-GFP* in**(E)** and **(F)**. The insets in the lower right indicate the egg cell (arrows), zygotes or embryos (arrowheads). Note that at 3 DAP, the *pZOU::H2B-GFP* reporter expression is largely confined to the embryo-surrounding region (ESR) in both normal and embryo-free seeds. Scale bars, 20 µm.

As in many angiosperms, *Arabidopsis thaliana* endosperm development consists of two main phases: an initial syncytial phase followed by a cellularized phase. In the embryo-free seeds, auto-fluorescence analysis revealed that the initiation and progression of endosperm cellularization occurred normally (Figures 4A and 4C; Supplemental Figure 1B). Endosperm cell walls were present in the embryo-free seeds, as in WT seeds at 6 days after pollination (DAP), indicating that the initiation and progress of endosperm cellularization are independent of embryo–endosperm communication, more like an autonomous developmental process. AGL62, a Type I MADS domain protein (Kang et al., 2007), functions as a major negative regulator of endosperm cellularization in *Arabidopsis* and is exclusively expressed during the syncytial phase and then declines abruptly just before cellularization (Figure 4B). Interestingly, when the embryo was absent, the expression of *AGL62-GFP* was identical to that in the WT (Figures 4B and 4D). GFP signals were detectable at 3 DAP, but not at 5 DAP, indicating the disappearance of *AGL62* expression according to the normal programmed time schedule, confirming normal endosperm cellularization in embryo-free seeds.

**Figure 4.**
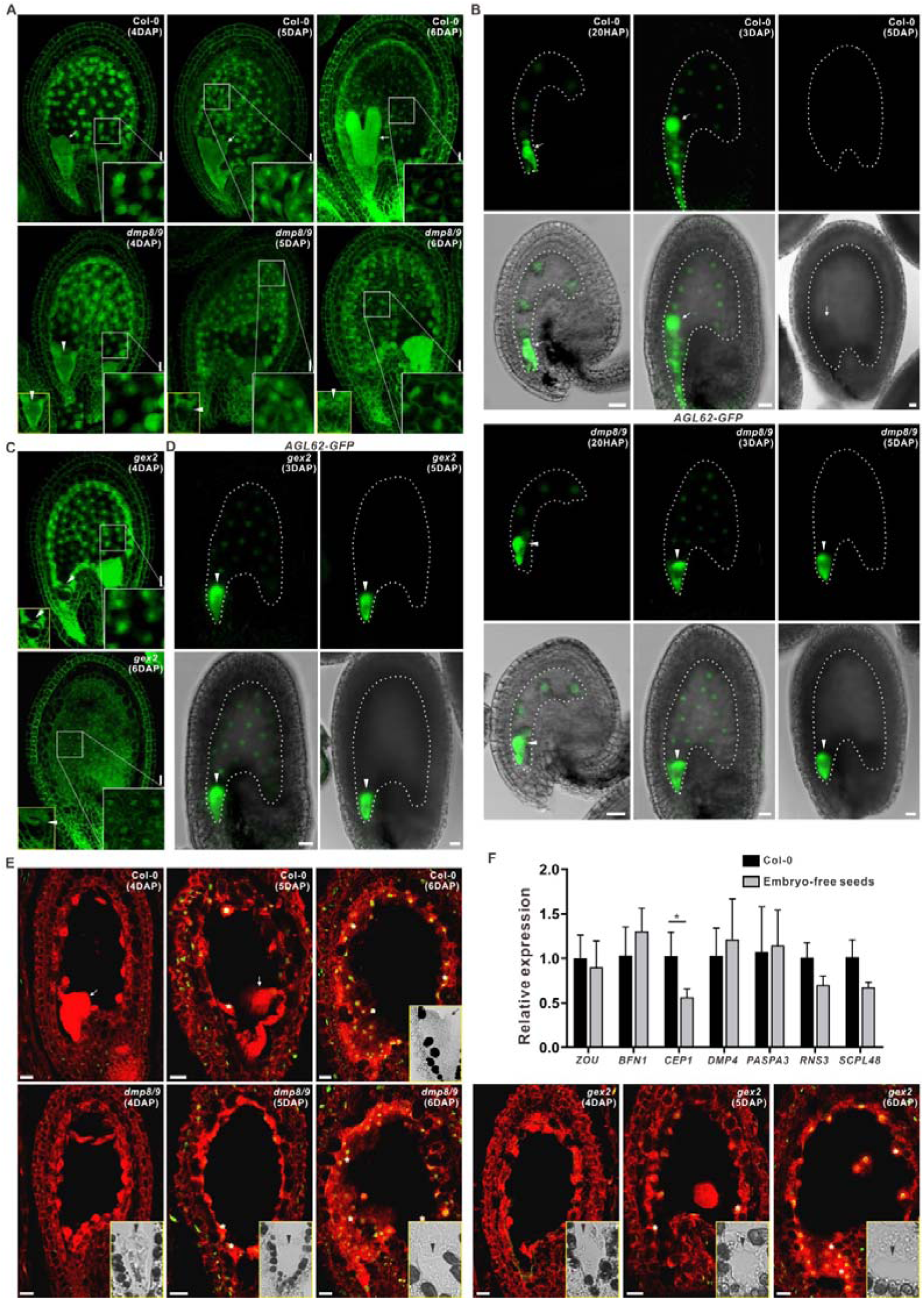
Endosperm cellularization is a cell-autonomous process and the embryo is not required for the initiation of endosperm cell PCD. **(A)** Autofluorescence of seeds seen by confocal microscopy after pollination with Col-0 or *dmp8/9* pollen. The insets in the lower left indicate the egg cell. **(B)** Seeds expressing *pAGL62::AGL62-GFP* after pollination with Col-0 or *dmp8/9* pollen. Note that *AGL62*, which suppresses cellularization, is expressed during syncytial endosperm development and becomes undetectable before cellularization in both normal and embryo-free seeds. The egg cell and embryo are marked by *pDD45::GFP*. **(C)** Autofluorescence-analysis of endosperm cellularization in embryo-free seeds pollinated with *gex2* pollen. The insets in the lower left indicate the egg cell. **(D)** Embryo-free seeds pollinated with *gex2* pollen show a *pAGL62::AGL62-GFP* expression pattern similar to that of Col-0 seeds. The egg cell is marked with *pDD45::GFP*. TUNEL signals in seeds pollinated with Col-0, *dmp8/9*, or *gex2* pollen. Propidium iodide staining was used to stain cells and the **(E)** TUNEL-positive signal is indicated by asterisks, and shown by the yellow fluorescence (green + red). Note that PCD signals begin to appear at 5 DAP in embryo-free seeds, similar to the control. Arrows indicate the embryo; arrowheads indicate the egg cell. Scale bars, 20 µm in **(A–E).** **(F)** qRT-PCR analysis of *ZOU* and developmental PCD markers in embryo-free seeds compared with the WT (Col-0) at 6 DAP. Data are the means ± SD of three biological replicates. Significant differences (\**P* < 0.05, two-sided Student’s *t*-test) are indicated.

In *Arabidopsis*, after cellularization, the endosperm eventually experiences cell death and is gradually absorbed by the embryo, which lives on to form the plant of a new sporophyte generation. Previous work using *dek1-3* and *atml1-3 pdf2-2* mutants reported that endosperm breakdown requires embryo growth in which embryo development arrests at the globular stage, and then the endosperm remains intact (Fourquin et al., 2016). Here, in embryo-free seeds, which completely exclude the influence of embryo growth and the risk of gene expression leakage, we found that the endosperm cell wall was still present at 9 DAP, as previously reported, while in phenotypically WT seeds, the endosperm had been eliminated (Supplemental Figure 2A). This means that rapid endosperm breakdown indeed involves embryo growth in *Arabidopsis*. To understand whether the initiation of endosperm programmed cell death (PCD) relies on embryo development due to spatial competition or embryo-derived signals, we used terminal deoxynucleotidyl transferase dUTP nick end labeling (TUNEL) to follow DNA degradation *in situ*, which highlights ongoing PCD in the endosperm cells. In the WT, TUNEL signals began to appear in endosperm cells at 5 DAP and became widespread at 6 DAP (Figure 4E). Surprisingly, although the embryo was absent, the endosperm showed the same TUNEL signal patterns, a sign of the initiation of endosperm degeneration, suggesting the embryo-independent initiation of DNA fragmentation. Although the mechanism underlying endosperm breakdown remains unclear, *ZOU* was reported to be responsible for endosperm breakdown (Yang et al., 2008) and triggered cell death by regulating the expression of cell-wall-modifying enzymes (Fourquin et al., 2016). Studies have also demonstrated that different cases of developmental PCD share a set of cell death-associated genes (Olvera-Carrillo et al., 2015). Therefore, we characterized the expression of *ZOU* and six of these potential PCD markers in the endosperm (Supplemental Figure 2B) during seed development using qRT-PCR (Figure 4F). This showed that except for *CEP1*, the expression of *ZOU* and the other five developmental PCD markers is not altered in embryo-free seeds compared with the control, confirming the embryo-independent initiation of endosperm breakdown.

Previous work has demonstrated that co-ordination between the endosperm and the seed coat is necessary for successful seed development (Roszak and Kohler, 2011; Garcia et al., 2003; Wang et al., 2010; Luo et al., 2005) and thus we also observed the process of seed coat development in different fertilization types of seeds when pollinated with *dmp8/9* pollen (Figure 5). In seeds containing only an embryo, seed coat development was not initiated and the integument began to degenerate at 4DAP, similar to that of unfertilized ovules (Figures 5C to 5E). This finding is consistent with previously reported results that the embryo alone is not sufficient for the initiation of seed coat growth (Roszak and Kohler, 2011). In *Arabidopsis*, upon double fertilization, the surrounding integuments will undergo a process of growth and differentiation that will lead to the formation of five cell-layered seed coat: two layers derived from the outer integument and three layers derived from the inner integument. Obviously, the seed coat in all embryo-free seeds was well initiated and normally developed. The seed coat was composed of five layers, the same as that in WT seeds (Figures 5A, 5B, 5E and 5F), suggesting that seed coat development mainly depends on endosperm development and conversely confirming normal endosperm development when the embryo is absent. In fact, we could not morphologically distinguish the embryo-free seeds from WT seeds until 6 DAP. Later, the embryo-free seeds showed smaller in size (Figures 5G; Supplemental Figures 3B and 3C), indicating that when endosperm PCD is initiated, the embryo plays an increasingly important role in seed expansion and further development, revealing an interesting coordination between seed coat and a late embryo although the molecular signaling between them remains to be elucidated. In *Arabidopsis*, seed coat growth is mostly driven by cell elongation (Garcia et al., 2005) and this was also illustrated by our findings that the cell number of the outermost seed coat layer is not changed but the average cell length is increased during the progress of seed coat development, no matter in normal seeds or embryo-free seeds (Supplemental Figure 3A).

**Figure 5.**
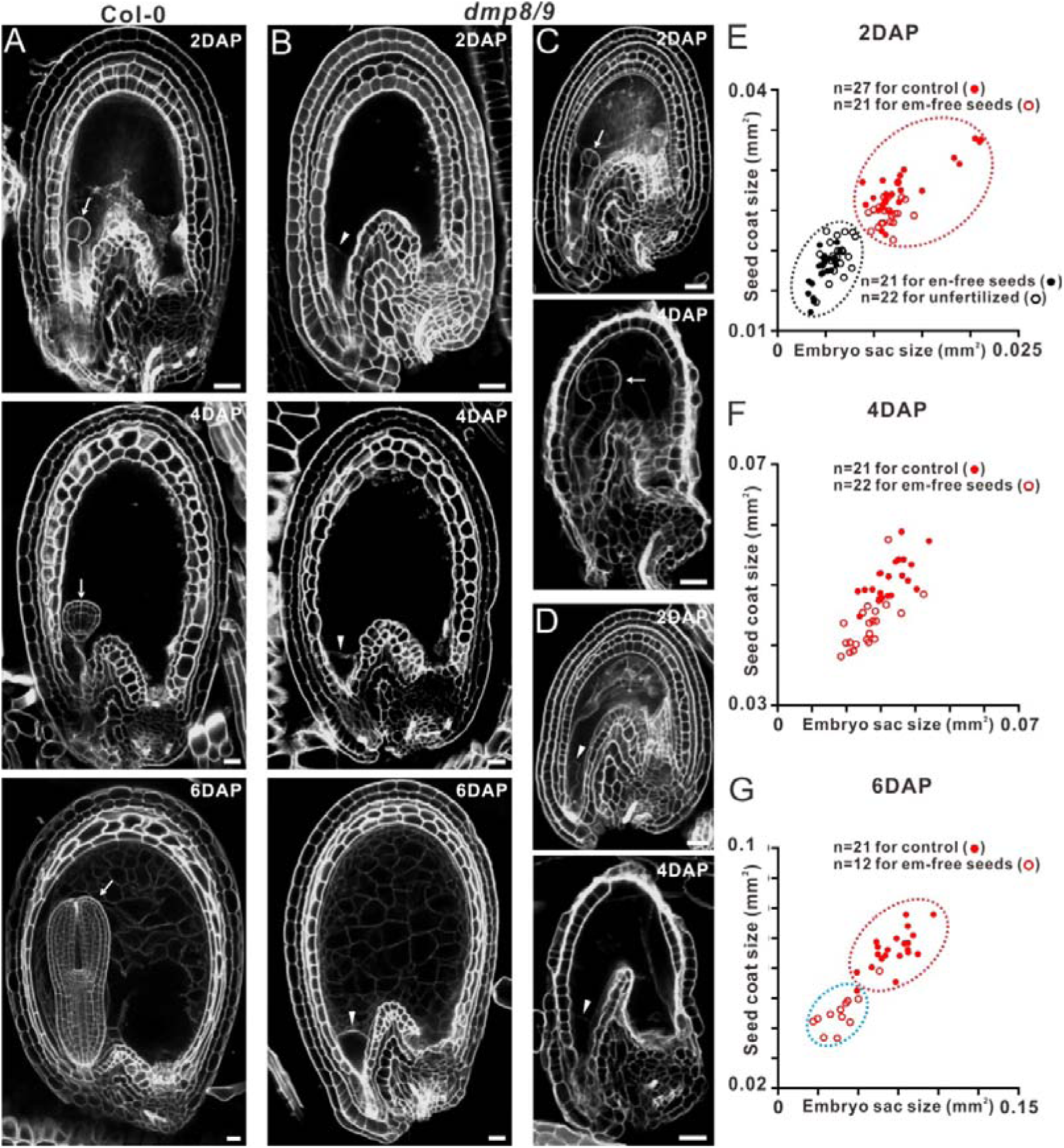
Seed coat development in embryo-free seeds. **(A)** Propidium iodide-stained seeds in WT at 2, 4 and 6 DAP. **(B-D)** Propidium iodide-stained seeds after pollination with *dmp8/9* pollen at 2, 4 and 6 DAP. Please note that the seed coat development in embryo-free (em-free) seeds **(B)** appeared to be similar to normal seeds **(A)**. Meanwhile seed coat development in seeds containing no endosperm (en-free) was not initiated and the integuments collapsed as that in unfertilized ovules (un-fer) at 4 DAP **(C-D)**. Arrows indicate the embryo and arrowheads indicate the egg cell. Scale bars, 20 µm. **(E-G)** The distribution of seed coat and embryo sac sizes in seeds after pollination with Col-0 or *dmp8/9* pollen at 2, 4 and 6 DAP, respectively.

In summary, our results strongly support the conclusion that the embryo is not required for endosperm development, although embryo growth acts in rapid endosperm elimination. In *Arabidopsis*, the endosperm is an ephemeral tissue that breaks down almost completely to provide space for embryo expansion physically and to recycle the nutrients stored in the endosperm tissues to fuel embryo growth. When an embryo is not present, endosperm elimination seems meaningless. Surprisingly as we observed, endosperm cells begin PCD as usual and ultimately break down almost completely at a low speed (Figure 4E and Supplemental Figure 2C). Thus, endosperm development is actually an autonomously programmed process, independent of embryo development. This work provides direct evidence for an “altruistic” nature of the endosperm in the relationship with its “sibling”-the zygotic embryo, and also a self-directed role in embryo-endosperm coordinated development.

Recently, auxin has been reported to be a signal involved in the dialogue among the endosperm, embryo, and seed coat. In *Arabidopsis*, auxin production after fertilization in the central cell is sufficient to trigger endosperm proliferation (Figueiredo et al., 2015; Batista et al., 2019). Intriguingly, auxin effIux from the endosperm has been reported to drive seed coat development (Figueiredo et al., 2016). Furthermore, auxin derived from the integument appears to be required for correct embryo development (Robert et al., 2018). Although direct links between endosperm-derived auxin and embryo development remain elusive, current knowledge suggests a central controller role of the endosperm in seed development. Thus, it is not surprising that the endosperm self-programs all its critical developmental processes and promotes seed coat development. In addition, considering the origin of double fertilization, as seen in *Ephedra*, sperm cells fuse with identical female nuclei to produce an embryo and a supernumerary embryo (Friedman et al., 1998). Both embryos develop independently and the development of the second fertilization product is characterized by an initial period of free nuclear proliferation followed by a process of cellularization, just similar to that of endosperm in flower plants. Thus, our finding seems favor the homologous theory, which can well explain the independence of endosperm development.

## METHODS

### Plant materials and growth conditions

*Arabidopsis thaliana* (accession Col-0) were grown in greenhouse under a photoperiod of 16 h light and 8 h dark at 22°C. The T-DNA insertion lines *gex2-2* (FLAG_441D08) (Mori et al., 2014) was obtained from the Nottingham Arabidopsis Stock Centre (NASC). The background of all Arabidopsis marker lines was Col-0.

### Constructs and plant transformation

For the construct of *ProZOU::H2B-GFP*, a 1.5kb promoter was amplified from genomic DNA using the primer pair (*ZOU-H2B-S/A*) and cloned into destination of the vector pART27 upstream of H2B-GFP after the digest of KpnI and AvrII. For the double marker lines carrying *pDD45::GFP* and *pDD22::CFP*, the length of promoter used in this study is according to the previous report (Steffen et al., 2007). For the *DMP8/9* CRISPR-Cas9 vector, *DMP8* and *DMP9* were targeted by one sgRNA and generated using a robust CRISPR/Cas9 vector system according to the reported methods (Ma et al., 2015).

All constructs were verified by sequencing and subsequently transformed into *Arabidopsis* (Col-0) by floral dip methods (Clough and Bent, 1998). Gene editing events for *DMP8/9* were analyzed by amplifying the genomic region that flanks the sgRNA target site by PCR, followed by sequencing. *dmp8/9-1* was used for subsequent experiments.

### Cytological observation

Ovule clearing was performed as previously reported (Boisnard-Lorig et al., 2001) and ovule autofluorescence observation was used to analysis endosperm cellularization (Li et al., 2017). As for TUNEL assays, a reported method was adapted using the DeadEndTM Fluorometric TUNEL System (Promega) (Wang et al., 2019). For Propidium iodide-staining seeds, Schiff reagent with propidium iodide (P4170; Sigma) was used like previously described (Shi et al., 2019) and finally the samples were observed under a confocal microscope (Leica SP8 CLSM). The area and length was calculated using the “measure”-tool with ImageJ.

### RT-qPCR

At 6 DAP, the ovules which smaller than siblings side by side were dissected from siliques when *dmp8/9-1* as a pollinator. Totally RNA of about 250 ovules a sample was extracted using RNeasy Plant Mini Kit. After digestion with DNase I (Qiagen), first-strand cDNA synthesis was performed using an M-MLV First-Strand Kit (Invitrogen, Carlsbad, CA, USA). RT-qPCR analysis was conducted according to the protocol previously described (Czechowski et al., 2005). The data were normalized to three housekeeping genes (*AT4G0532, AT1G13320* and *AT4G34270*), and each experiment was repeated three times.

### Accession Numbers

The Arabidopsis Genome Initiative accession numbers for the genes and gene products mentioned in this article are as follows: At1g09157 (*DMP8*), At5g39650 (*DMP9*), AT5G49150 (*GEX2*), At2g06090 (*DD19*), At5g38330 (*DD22*), At5g48670 (*AGL80*), AT5G60440 (*AGL62*), AT2G15890 (*AtCBP1*), AT1G49770 (*ZOU*), AT4G04460 (*PASPA3*), AT1G11190 (*BFN1*), AT4G18425 (*DMP4*), AT5G50260 (*CEP1*), At1g26820 (*RNS3*) and At3g45010 (*SCPL48*).

## ACKNOWLEDGMENTS

We thank Gary Drews for kindly providing maker lines (*pDD19::GFP, AGL62-GFP* and *AGL80-GFP*) and Wei-Cai Yang for the *CBP1-3xGFP* maker line. We thank Y.G. Liu for providing the CRISPR/Cas9 system used in this work. This work was supported by two grants from National Natural Science Foundation of China (31991201; 31800264).

## AUTHOR CONTRIBUTIONS

H.X., W.W., and MX.S. designed the experiments. HX.X. and W.W. performed the experiments. W.W, H.X. and MX.S. analyzed the data and wrote the manuscript. All authors approved the final version of the manuscript.

## Supplemental Data

**Supplemental Figure 1.**
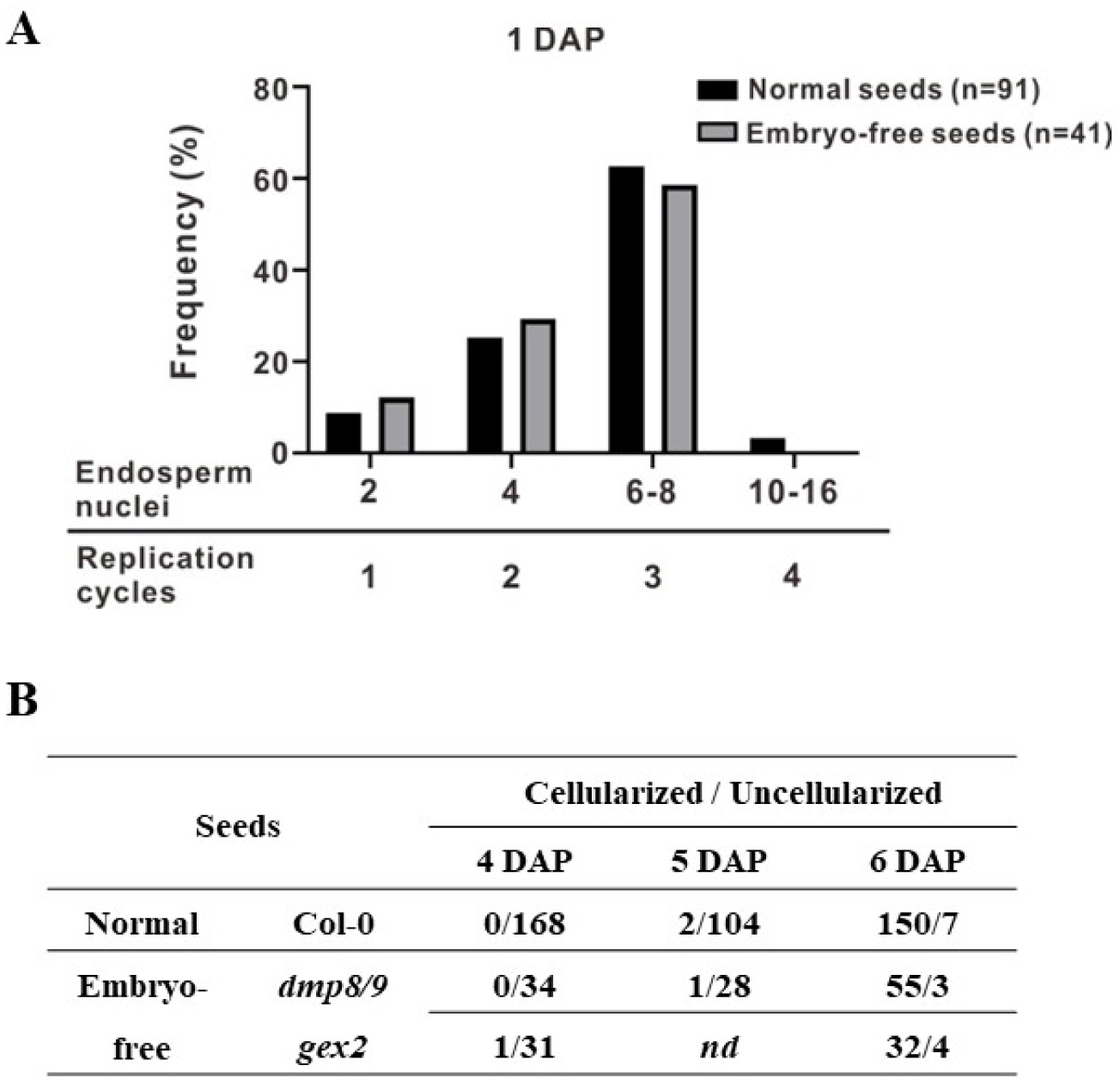
Phenotype analysis of endosperm in Col-0 and embryo-free seeds. **(A)** Distribution of endosperm nuclei number in Col-0 and embryo-free seeds shown in Figure 3A. At 1 DAP, embryo-free seeds pollinated with *dmp8/9* pollen do not have defects in endosperm proliferation. **(B)** Statistics of endosperm cellularization in seeds at 4-6 DAP.

**Supplemental Figure 2.**
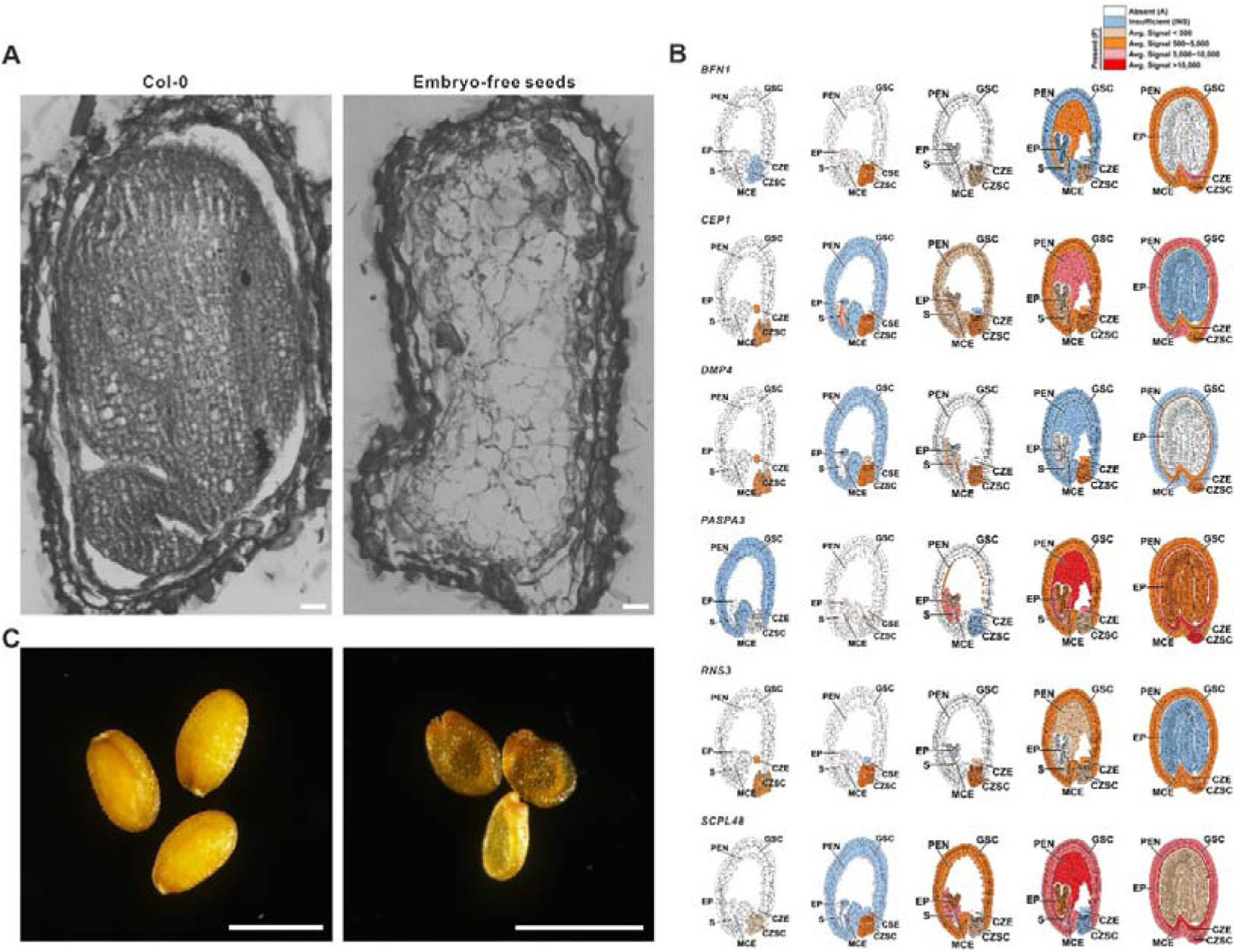
Embryo growth accelerates endosperm breakdown. **(A)** Toluidine Blue-stained paraffin sections showing endosperm structure at 9 DAP. Please note that in wild-type seeds, the endosperm had been eliminated. However, in embryo-free seeds, the endosperm was intact. Scale bars, 20 µm. **(B)** Expression data for *BFN1, CEP1, DMP4, PASPA3, RNS3* and *SCPL48* downloaded from the Seed Gene Network resource (http://seedgenenetwork.net/). **(C)** Dry seeds of Col-0 and embryo-free seeds in *dmp8/9*. Scale bars, 0.5 mm.

**Supplemental Figure 3.**
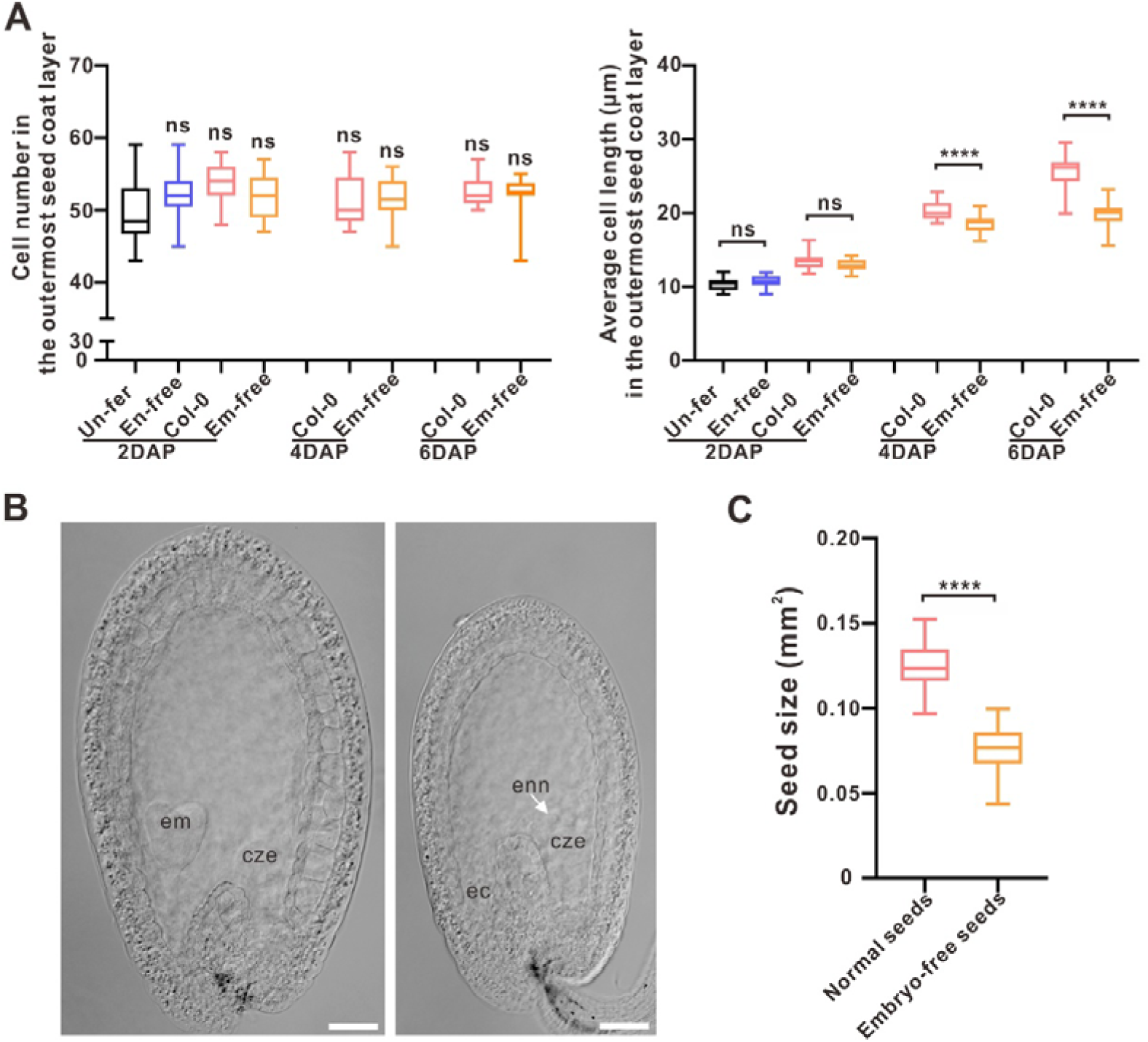
Integument cell elongation is responsible for seed coat growth and embryo-free seeds show smaller sizes at 6 DAP. **(A)** Quantification of cell number and length in the outermost seed coat layer shown in Figure 5. **(B)** Cleared *dmp8/9* ovules, 6 days after pollination. Developing embryo and endosperm (left); endosperm but no embryo (right). Scale bars =50 µm. **(C)** Quantification of seed sizes shown in **(B)**, indicating that the embryo-free seeds are smaller. Data are the means ± SD, with n= 38, 48 seeds from left to right. Significant differences (**** P < 0.0001, two-sided Student’s t-test) are indicated. Abbreviations: un-fer: un-fertilized; en-free: endosperm-free; em-free: embryo-free; em: embryo; ec: egg cell; enn: endosperm nuclei; cze: the chalazal endosperm.

**Supplemental Table 1.**
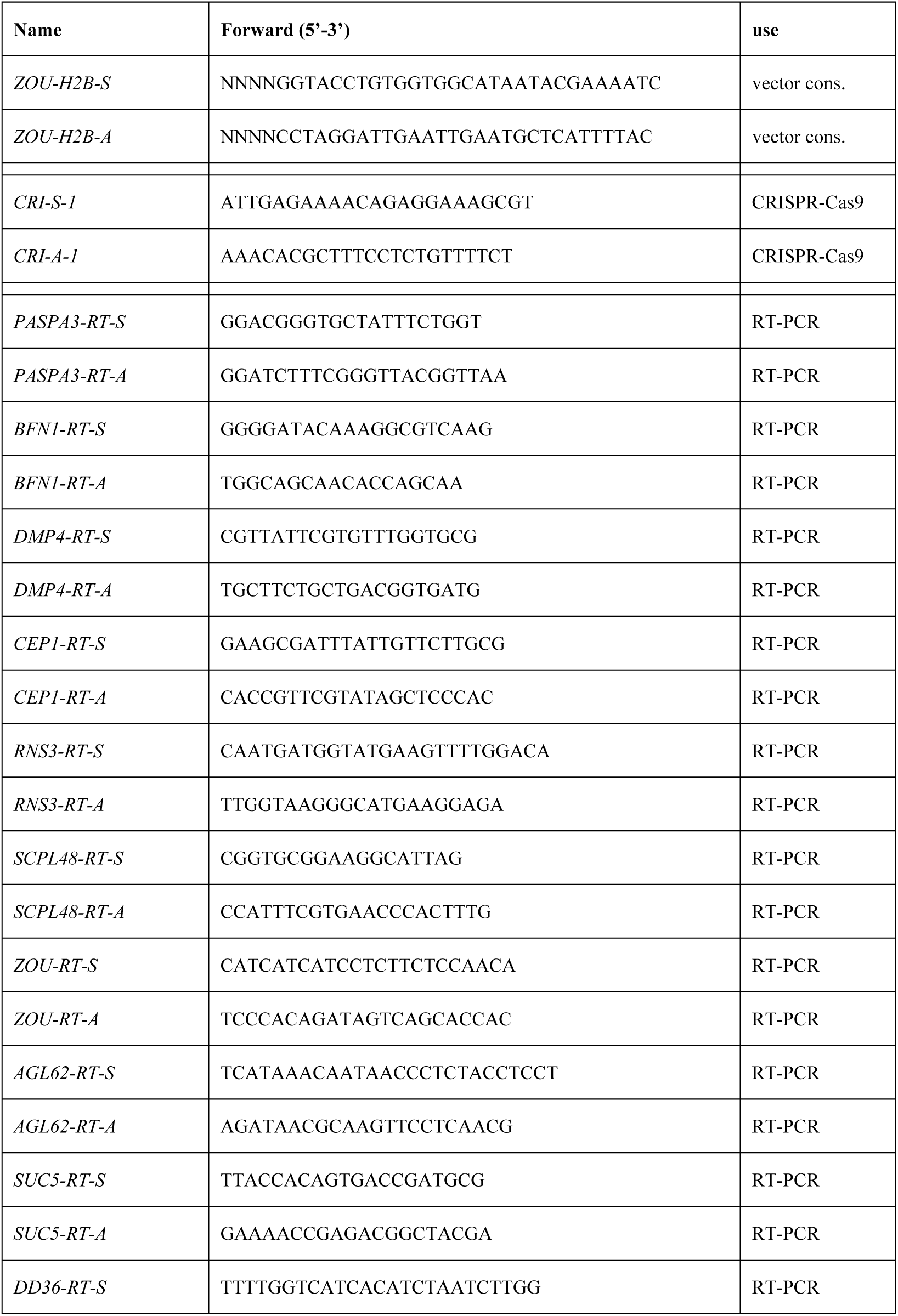
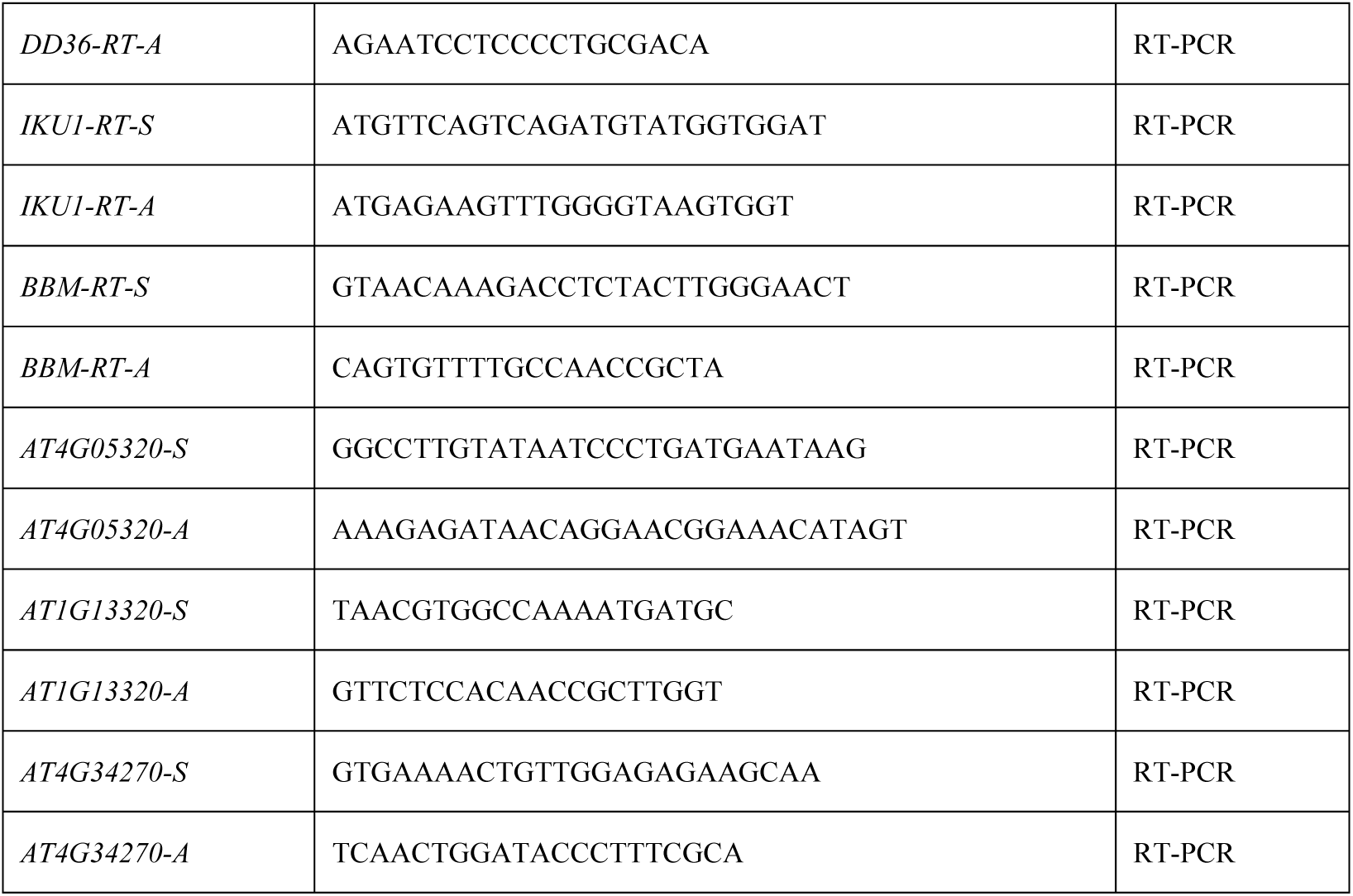
Primers used in this study

